# Optogenetic-Induced Muscle Loading Leads to Mechanical Adaptation of the Achilles Tendon Enthesis in Mice

**DOI:** 10.1101/2023.04.11.536376

**Authors:** Elahe Ganji, Syeda N. Lamia, Matthew Stepanovich, Noelle Whyte, Adam C. Abraham, Megan L. Killian

## Abstract

The growth of the skeleton depends on the transmission of contractile muscle forces from tendon to bone across the extracellular matrix-rich enthesis. Loss of muscle loading leads to significant impairments in enthesis development. However, little is known about how the enthesis responds to increased loading during postnatal growth. To study the cellular and matrix adaptations of the enthesis in response to increased muscle loading, we used optogenetics to induce skeletal muscle contraction and unilaterally load the Achilles tendon and enthesis in young (i.e., during growth) and adult (i.e., mature) mice. In young mice, daily bouts of unilateral optogenetic loading led to expansion of the calcaneal apophysis and growth plate, as well as increased vascularization of the normally avascular enthesis. Daily loading bouts, delivered for 3 weeks, also led to a mechanically weaker enthesis with increased molecular-level accumulation of collagen damage in young mice. However, adult mice did not exhibit impaired mechanical properties or noticeable structural adaptations to the enthesis. We then focused on the transcriptional response of the young tendon and bone following optogenetic-induced loading. After 1 or 2 weeks of loading, we identified, in tendon, transcriptional activation of canonical pathways related to glucose metabolism (glycolysis) and inhibited pathways associated with cytoskeletal remodeling (e.g., RHOA and CREB signaling). In bone, we identified activation of inflammatory signaling (e.g., NFkB and STAT3 signaling) and inhibition of ERK/MAPK and PTEN signaling. Thus, we have demonstrated the utility of optogenetic-induced skeletal muscle contraction to elicit structural, functional, and molecular adaptation of the enthesis *in vivo* especially during growth.

## Introduction

The enthesis is a tissue interface that adapts to applied muscle loads during postnatal growth by forming a graded fibrocartilage transition between tendon and bone (Rossetti et al. 2017; Michael Benjamin and McGonagle 2009; Thomopoulos et al. 2006; Golman et al. 2021). The mature enthesis is a network of mineralized and unmineralized collagen fibers that is tougher than either tendon or bone (Genin and Thomopoulos 2017). As the result of this structural toughening, acute and high energy-induced tendon injuries typically result in failure of tendon (e.g., Achilles tendon ruptures) or bone (e.g., calcaneal avulsion fractures) and not the enthesis (Shamrock and Varacallo 2021; Weatherall, Mroczek, and Tejwani 2010). However, with chronic loading, the enthesis is susceptible to localized damage and overuse-related injuries, and up to 25% of all tendon and ligament disorders involve the enthesis (Ogden et al. 2004; Duong et al. 2020; M. Benjamin et al. 2007; Maffulli, Wong, and Almekinders 2003). In fact, overuse is considered the underlying cause of most entheseal disorders (e.g., Achilles insertional tendinopathy, Sever disease) that are most commonly observed in young athletes (Shopfner and Coin 1966; Ross and Caffey 1957; Ogden et al. 2004; Kvist 1991).

Yet loading is essential for functional development and growth of the tendon-bone enthesis (Schwartz et al. 2013; M. Benjamin et al. 2006; Shaw and Benjamin 2007; Zelzer et al. 2014; Shwartz, Blitz, and Zelzer 2013; Felsenthal and Zelzer 2017; Killian 2022). During embryonic development, tendon extends from the cartilage anlagen to guide load transfer from skeletal muscle which induces the mechano-adaptive growth of the enthesis (Blitz et al. 2009). During postnatal development, the enthesis matures into a graded fibrocartilage interface from the fibrous tendon to the mineralized bone (Schwartz et al. 2013; M. Benjamin et al. 2006). The structural adaptation of the enthesis occurs in response to a combination of tensile and compressive loads (Shaw and Benjamin 2007; M. Benjamin et al. 2006; Golman et al. 2021). While several elegant studies have established the requirement of muscle contraction during musculoskeletal growth, these studies have primarily focused on the absence or reduction of muscle loading rather than increased muscle contraction (Blitz et al. 2009; Kahn et al. 2009; Rot-Nikcevic et al. 2006; Dysart, Harkness, and Herbison 1989; Ford et al. 2017; Giorgi et al. 2015). Increased activity during periods of rapid growth, such as in children and during adolescence, can lead to micromotion at the enthesis and apophysis and result in pain and clinical presentation of disorders such as Sever disease (Duong et al. 2020; Ogden et al. 2004). In adults, repeated tendon loading during activities like running-associated overuse can lead to abnormalities such as bony spur formation at the distal tendon that leave individuals with pain and dysfunction (M. Benjamin et al. 2006; Michael Benjamin and McGonagle 2009; M. Benjamin et al. 2007; 2009). Despite the potential age-associated differences in adaptation of the enthesis to repeated loading, the mechanisms contributing to enthesis remodeling in the immature and mature skeleton remain poorly understood and challenging to study (Achar and Yamanaka 2019).

In mouse and rat studies of skeletal adaptation to mechanical loading, cyclic, sub-failure compressive loading has commonly been used to induce anabolic bone formation in both the ulna and tibia (Lynch et al. 2010; De Souza et al. 2005; Fritton et al. 2005; Patel, Brodt, and Silva 2014; Kotha et al. 2004; Robling et al. 2002). Similarly, sub-failure cyclic tensile loading of the patellar tendon has been used to model tendon fatigue and damage in mice and rats (Fung et al. 2010a; Andarawis-Puri et al. 2012; Sereysky et al. 2012). Unfortunately, the latter approach does not load tendon enthesis via muscle contraction but instead via direct, open, and external mechanical actuation. More commonly used and physiologically-relevant models of tendon loading, such as forced treadmill running (uphill or downhill), are effective for inducing structural changes to the mature enthesis in response to increased loading but also have their drawbacks (Soslowsky et al. 2000; Abraham et al. 2019; Thampatty and Wang 2018). For example, treadmill running studies in mice and rats, like most bone anabolic loading studies, typically initiate forced running at 7-8 weeks of age (Kim et al. 2020), which limits our ability to study tendon and enthesis adaptation during maturation. Additionally, these approaches are highly variable, as even inbred mouse strains have high variations in distances run and tolerance for forced treadmill running (Lerman et al. 2002). Forced treadmill running can also confound results by inducing anxiety, leading to systemic inflammation (M. Svensson et al. 2016) as well as effect organismal metabolic conditioning induced by cardiovascular load and training effects (Koch et al. 2011). To counter the systemic effects and reduce loading dose variability, electrical nerve stimulation has also been used to induce repeated muscle stimulation in mice and rats for tendon loading (Rezvani et al. 2021; Nakama et al. 2005). However, this approach is also challenging, as most studies are also in adult animals and these approaches require subdermal needle electrode placement which may increase risk of injury and infection with repeated electrode placement in the growing muscle and tendon. Thus, understanding how loading effects structural changes to the tendon enthesis during growth has been challenging *in vivo* because of a lack of physiologically relevant loading models.

To address these challenges, we turned to the use of optogenetics as a non-invasive tool for direct and precise induction of muscle contraction for *in vivo* tendon loading in growing mice. Optogenetics is a tool commonly used in neuroscience that facilitates controlled activation of activatable cells such as neurons and muscle cells, with high spatial and temporal specificity (Boyden et al. 2005; Boyden 2011). Channelrhodopsin-2 (ChR2) is a light-sensitive microbial opsin that, when present on the cell membrane, acts as a non-specific light-activated cation channel that induces membrane depolarization (Nagel et al. 2003; Deisseroth 2011). Optogenetic stimulation improves the spatial specificity of electrical stimulation by targeting cell subpopulations based on morphological markers and molecular footprint (Boyden et al. 2005; Boyden 2011; Jia et al. 2011; Bruegmann et al. 2010; Magown et al. 2015). We and others have previously shown that optogenetic-mediated activation of skeletal muscle allows for precise and controlled contraction of specific muscle groups (Magown et al. 2015; Ganji et al. 2021; Bruegmann et al. 2015) and, therefore, their associated tendons and entheses. In this study, we examined how sustained muscle contractions, induced by optogenetic activation, can differentially elicit structural, functional, and molecular changes in the young and adult Achilles enthesis of mice. We then defined transcriptional changes in the Young tendon and bone following repeated loading.

## Results

### Optogenetic activation is a repeatable and safe approach for daily tendon loading

To investigate the effects of increased loading on the postnatal Achilles enthesis, we generated a transgenic mouse line to express ChR2 in skeletal muscle using doxycycline-treated Ai32-floxed, ACTA1-rtTA;tetO-Cre mice (Ganji et al. 2021) and unilaterally loaded their Achilles tendon using optogenetic stimulation of the triceps surae muscle. Daily bouts of loading were delivered at either 5- or 20-minute bouts for 5 days a week, up to 3 weeks in duration, in mice starting at either 4 weeks of age (Young group; this time point coincides with enthesis growth) or at 6-8 months of age (Adult group, 20-min bouts only; mature enthesis). In Young mice, the steady state generated torque at the end of the 20-minute loading bout was ∼30% of the initial peak torque for the first week of daily optogenetic muscle loading. We found that daily loading in Young mice did not negatively impact animal weight gain, however some weight loss was observed in Adult mice (Figure 1A). Isometric ankle joint torque did not significantly vary over the course of the experiment for the Young group but did increase with daily 20-minute loading bouts in the Adult group from week 1 to week 2 of loading (Figure 1B).

**Figure 1.**
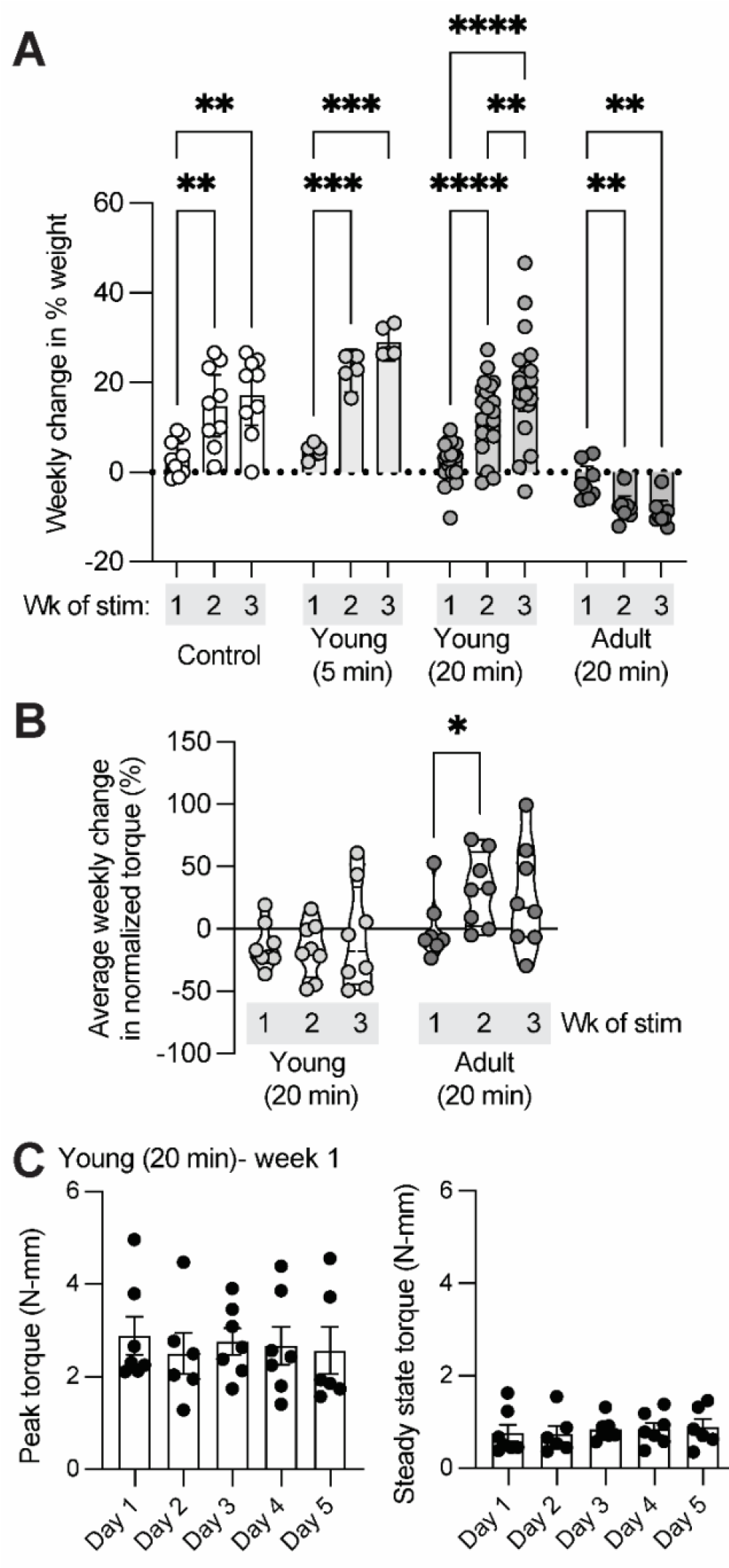
Optogenetic activation of triceps surae muscle group is not invasive and does not impact weight gain or generated isometric ankle joint torque in Young mice. (A) Young mice, regardless of exposure to daily bouts of optogenetic muscle stimulation (5- or 20-minute duration), gained weight for the duration of the experiment. However, adult mice lost weight from onset to end of the experiment. (B) Normalized torque significantly increased in adult but not young mice with daily bouts of 20-minute optogenetic muscle stimulation. Each data point denotes average weekly change in generated ankle torque normalized to weight of the animal. (C) The steady state generated torque at the end of the 20-minute loading bout, measured during isometric ankle plantarflexion, was ∼30% of the peak force in young mice for the first week of daily optogenetic muscle loading. Error bars: *Mean ± SD (*: p< 0*.*05, **: p< 0*.*001, ****: p < 0*.*0001, and ***: p = 0*.*0004)*.

### Daily tendon loading leads to bony changes and impaired mechanical properties of the Young but not Adult Achilles enthesis

Daily loading of the tendon and enthesis for three weeks led to apophyseal changes in both the Young and Adult calcanei (Figure 2A). More apparent changes were found in the Young group, with the calcaneus developing a “bump” and delayed fusion of the growth plate (Figure 2A). The Young loaded enthesis had increased vascular ingrowth compared to the contralateral enthesis, as visualized using histology (Figure 2B). In the Adult group, obvious surface shape changes were not observed, however three-dimensional reconstructions showed less mineral in the calcaneal apophysis (Figure 2A). This was evident for both Young and Adult calcanei following loading, which significantly reduced bone mineral density compared to the contralateral calcaneus (BMD; mg HA/cm^3^; Figure 2B; ****: p < 0.0001; ***: p = 0.0004). All tissues failed at the enthesis (Young and Adult) or calcaneal growth plate (Young), not the tendon mid-substance, during uniaxial testing. In Young mice, the maximum load before failure was significantly reduced following repeated loading compared to contralateral Achilles tendon and enthesis (Figure 2E). However, increased strength was observed in the Adult tendon/enthesis following 3 weeks of loading compared to the contralateral tendons (Figure 2E). For the Young, but not Adult, Achilles tendon/enthesis, toughness was significantly reduced after 3 weeks of loading compared to contralateral tendons (Figure 2E).

**Figure 2.**
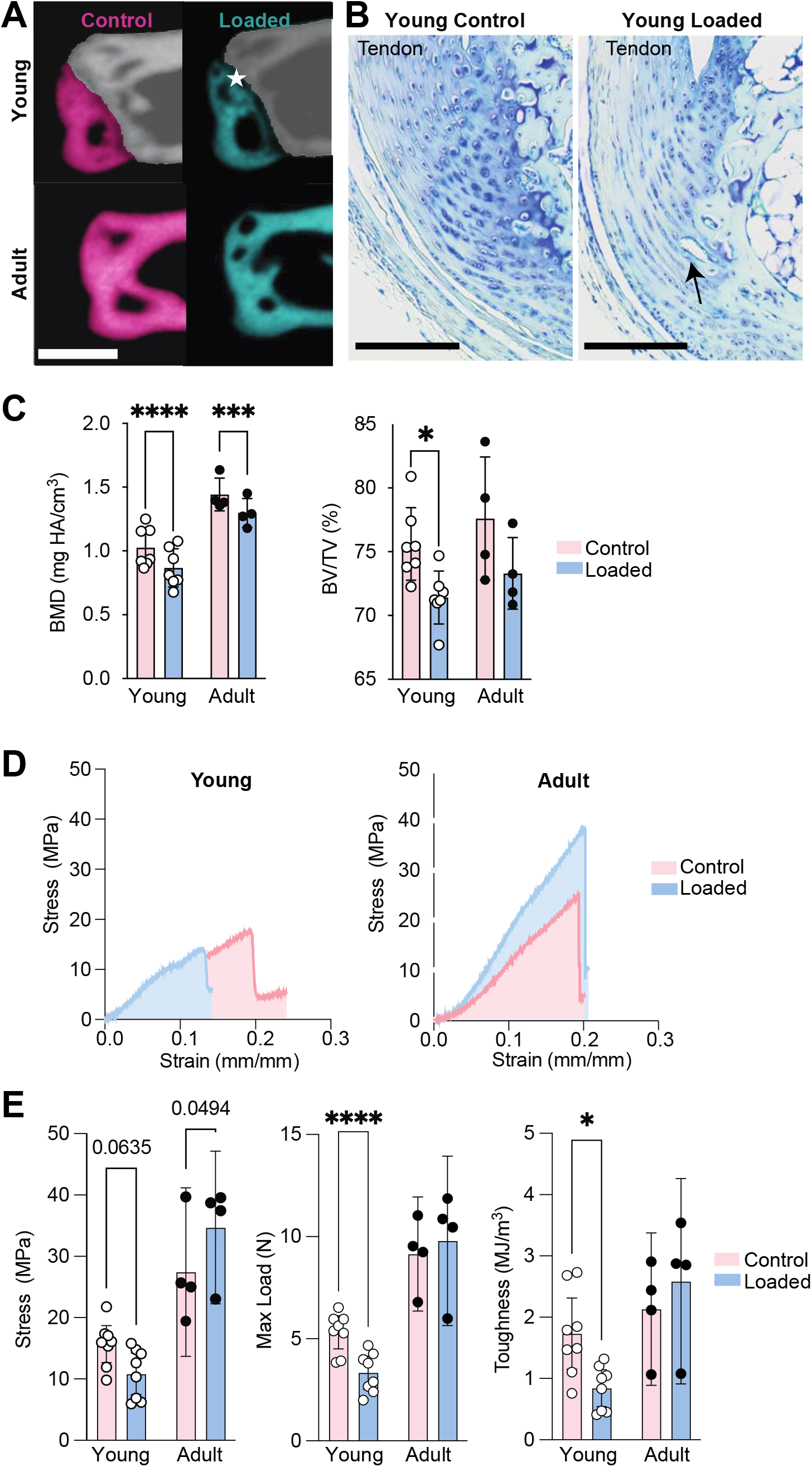
Repeated loading in Young mice results in disruption of structural toughening mechanisms at enthesis. Repeated loading resulted in (A) opening of the calcaneal growth plate, (B) disruption of enthesis tidemark, reduction of subchondral bone mineralization, and vascular infiltration, in Young enthesis and apophyses. (C) Young and Adult loaded apophyses had significantly lower BMD compared to age-matched controls. Structural adaptations in the Young enthesis and apophysis, coincides with (D, E) significantly less tensile strength and toughness in the loaded compared to non-loaded contralateral limbs. Star specifies the opening of the calcaneal growth plate. Scale bars denote (A) 500µm and (B) 100µm. (*: p< 0.0.5, **: p< 0.001, ****: p < 0.0001, and ***: p = 0.0004).

### Daily tendon loading leads to accumulated collagen damage in Young mice

Loading can initiate accumulative damage at the molecular level by inducing fibril breaks and protease-induced remodeling of the ECM. As a result of this ECM damage, collagen triple helices can irreversibly unfold (Zitnay et al. 2017). To evaluate collagen damage in our *in vivo* model of tendon loading, we used collagen hybridizing peptides (CHPs) conjugated with fluorescent labels to visualize molecular-level damage *in vivo* (Li et al. 2012; Bennink et al. 2018). In tendon, CHPs can specifically detect non-recoverable (post-yield) unfolding of collagen fibers (Zitnay et al. 2017). After 5 consecutive days of 20-minute daily loading bouts, Young loaded tendon and enthesis had higher CHP intensity (Figure 3B) and, therefore, collagen denaturation than contralateral controls (Figure 3C, p=0.0074). Collagen type III is correlated with increased remodeling, and loaded attachments had more Collagen type III and Aggrecan present at the enthesis compared to contralateral controls (Figure 3D). In addition, the calcaneal growth plate in loaded limbs had more Aggrecan than contralateral controls (Figure 3D).

**Figure 3.**
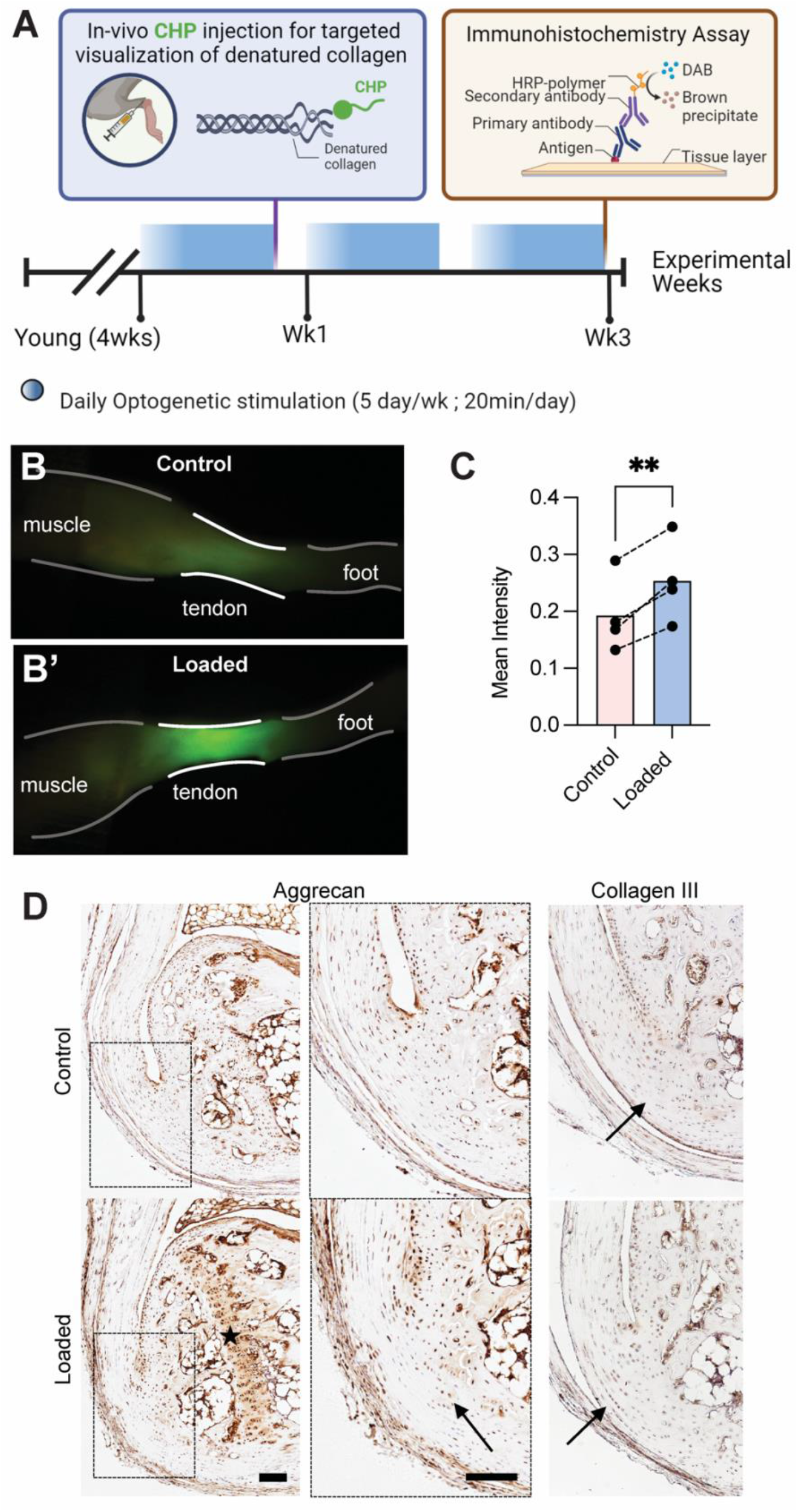
Repeated loading led to denatured collagen fibers and increased prevalence of ECM components at the enthesis and growth plate. (A) Schematic shows experimental design for evaluation of effect of repeated loading on ECM of the Young tendon and enthesis. (B, C) Repeated loading resulted in the accumulation of collagen denaturation in loaded tendon and enthesis. P<0.05. (D) Loaded entheses had qualitatively higher aggrecan and type III collagen present at the calcified and uncalcified fibrocartilaginous site of their attachments, respectively, compared to controls. Scale bar denotes 100 µm.

### Transcriptomics

We performed bulk RNA-seq with RNA isolated from Achilles tendons and calcanei of naïve and optogenetically-loaded mice (loaded and contralateral limbs) at 3hr post-final loading bout to investigate the molecular changes underlying the structural enthesis and bone morphology. We used principal component analyses (PCA) to evaluate data quality in an unsupervised approach (Figure 4A). To avoid any compensatory effects of the contralateral limb and because we saw discrete separation of naïve from loaded (5 day and 12 day) tendons in PCA, we used naïve data to measure fold change between optogenetically loaded and cage activity tendons and bones (Figure 4A). We identified 1,193 and 312 genes in tendon and bone, respectively, that were differentially expressed at both 5 and 12 days of loading (Figure 4B), and both tendon and bone had more differentially expressed genes at 12 days compared to 5 days of daily loading.

**Figure 4.**
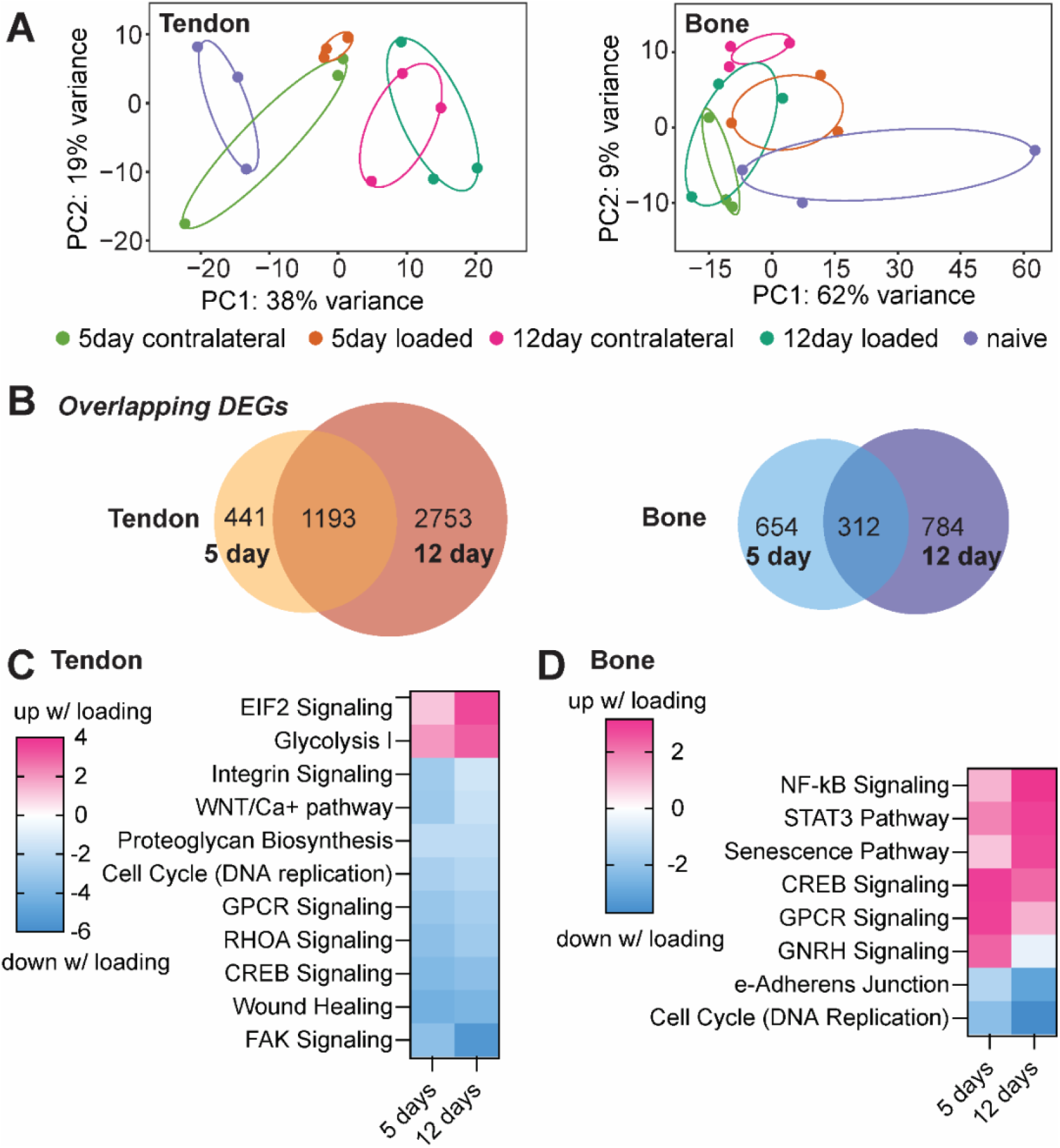
(A) Principal component analysis plots of differential gene expression for naïve, contralateral, and loaded tendons and bones after 5day and 12 days of loading. (B) In tendon and bone 1,193 and 312 genes, respectively, were differentially expressed after 5 and 12 days of loading. (C) In tendon, optogenetic loading induced upregulation of EIF2 signaling and Glycolysis I pathways, and downregulation of FAK and RHOA signaling pathways. (D) In bone, optogenetic loading led to activation of inflammatory pathways such as NFκB and STAT3 signaling, senescence, and CREB signaling, as well as downregulation of cell cycle (DNA replication) and epithelial adherens junction signaling.

We used Ingenuity Pathway Analysis (IPA, Qiagen, Germany) to predict upstream regulators and determine tissue-level loading-induced activated or inhibited pathways. In tendon, we found optogenetic loading for both 5 and 12 days led to activation of eukaryotic initiation factor 2 (EIF2) signaling and glycolysis (Figure 4C). Surprisingly, mechanical loading of tendon induced by optogenetic contraction of skeletal muscle resulted in downregulation of canonical pathways related to mechanobiology, including FAK and RHOA signaling (Figure 4C). In bone, loading led to activation of genes associated with inflammatory pathways such as NF-kB and STAT3 signaling as well as senescence and CREB signaling (Figure 4D). Loading also inhibited cell cycle (replication) in bone at both 5 and 12 days (Figure 4D).

We found optogenetic loading resulted in downregulation of numerous genes associated with tendon stem cells, including *Tppp3* and *Fbn1*, which were down at both 5 and 12 days, and *Tnc*, which was down at 5 days only. Of the top 20 downregulated genes (based on log2FC) in tendon following loading, most were associated with extracellular matrix synthesis at one or both time points, including *Matn3, Acan, Col8a1, Thbs1, Col9a2, Bglap, Mmp9*, and *Adamts19* (Figure 5). In bone, daily bouts of loading resulted in increased expression of *Il1ra2, Ccl21c, Ccl21a, Gpnmb*, and *Ccl19* and other pro-inflammatory markers at one or both time points (Figure 5). We also found that *Ihh* was one of the top 20 downregulated genes in bone at both 5 days with loading (Figure 5).

**Figure 5.**
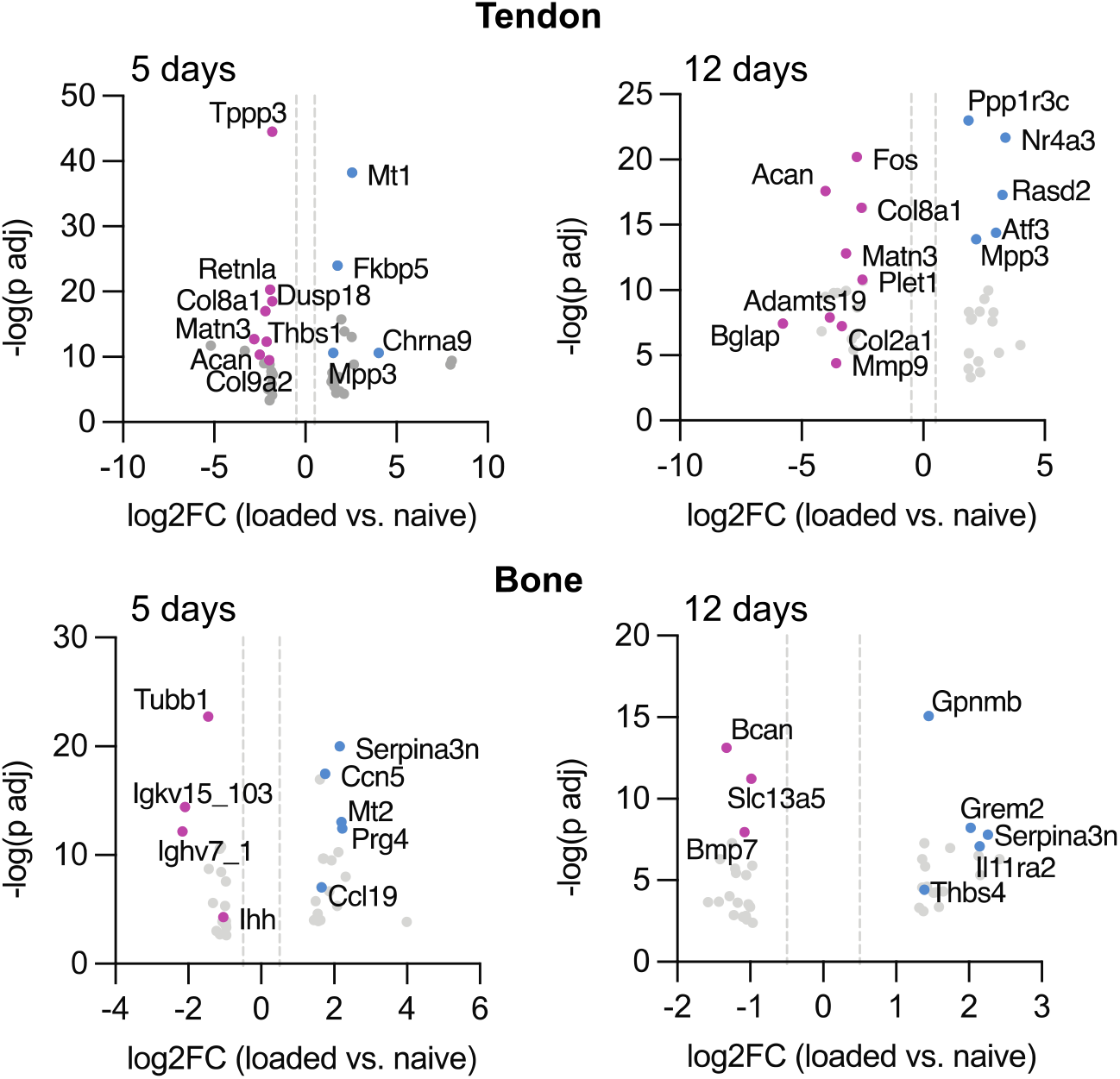
After 5 days and 12 days of stimulation, daily bouts of optogenetic loading led to significant downregulation of genes associated with tendon stem cell activation and ECM synthesis in tendon and upregulation of proinflammatory markers in bone. Blue indicates genes upregulated with loading compared to naïve and pink indicates genes downregulated with loading compared to naïve. Data shown are log2 fold change (x-axis) and -log10 transformed (y-axis).

## Discussion

In this study, we evaluated the mechanically induced adaptation of both the young and adult enthesis to repeated loading using optogenetic stimulation. Previous studies have explored the effects of repeated loading on the structural and functional adaptation of healthy and repaired tendon and enthesis in adult and aging tissues (Soslowsky et al. 2000; Lin et al. 2018; Cho et al. 2011; Pingel et al. 2013; Almeida-Silveira et al. 2000). Yet, the effect of repeated loading on immature tendon and enthesis was previously unknown. Repeated tensile loading of the Achilles tendon and enthesis using optogenetic activation of skeletal muscle was non-invasive and did not cause weight loss or gross macroscopic damage to the tendon or enthesis after 3 weeks. Repeated loading did result in microscopic remodeling and disruption of the structure and function of the young, but not the adult enthesis. Repeated loading also led to opening of the growth plate, increased vascular infiltration, disruption of the enthesis tidemark, and impaired mechanical properties in Young mice. This may have been caused by disruption of the continuous transition between the mineralized and unmineralized fibrocartilage of the enthesis that contributes to the tissue’s mechanical toughness (Hu et al. 2015; Liu et al. 2012) and is the primary determinant of enthesis mechanics and stress transfer between tendon and bone(Genin et al. 2009).

In tendons, aging leads to impaired self-renewal potential (Ruzzini et al. 2014; Kohler et al. 2013; Zhou et al. 2010; J. Zhang and Wang 2015), tissue turnover (R. B. Svensson et al. 2016), increased prevalence of proteoglycans and mineral (Sugiyama et al. 2019), and degenerative structural adaptations (e.g., reduced cellularity, disruption collagen fiber orientation, and fibroblastic changes) (Sugiyama et al. 2019; R. B. Svensson et al. 2016). In mice, reduced tendon stem cell proliferation has been reported as early as 5 months of age (J. Zhang and Wang 2015). In previous studies, degenerative changes of the adult tendon and enthesis in response to increased loading include increased cross-sectional area (Soslowsky et al. 2000), higher cellularity (Lin et al. 2018; Cho et al. 2011; Pingel et al. 2013), neovascularization (Lin et al. 2018), reduced fiber organization (Lin et al. 2018; Soslowsky et al. 2000; Pingel et al. 2013). Excessive repeated loading can also lead to reduced stiffness and strength (Almeida-Silveira et al. 2000). However, in our study, repeated muscle-induced loading of the adult enthesis did not lead to pathological adaptation, but improved structure and mechanics of the adult enthesis. This could be a result of physiological and isometric loading or post-loading compensatory recovery after dosing.

Increased and repeated loading leads to dose-dependent degeneration by the accumulation of sub-failure damage. Collagen is the primary stabilizing constituent of the tendon, bone, and enthesis ECM (Shoulders and Raines 2009) and the key contributing factor to the elastic behavior of tendon (Franchi et al. 2007) and toughness of the bone (Wang et al. 2001; Viguet-Carrin, Garnero, and Delmas 2006). In the tendon, structural reorganization of collagen fibers can be exacerbated into non-reversible molecular level denaturation of the collagen fiber collagen, ultimately leading to inferior mechanical properties (Zitnay et al. 2017; Fung et al. 2010b). Collagen hybridizing peptides (CHPs) are synthetic single strand peptides that have recently been developed to hybridize with denatured collagen triple helix. In tendon, CHPs can specifically bind to non-recoverable (post-yield) sites of collagen fibril unfolding (Zitnay et al. 2017). The increased *in vivo* labeling of the Achilles tendon with CHPs indicates that daily loading leads to increased collagen denaturation. Similar to findings from repeatedly loaded tendon *ex vivo* (Szczesny et al. 2018), our results show that loading-induced collagen denaturation does not coincide with altered tissue stiffness. These findings suggest a compensatory mechanism (e.g., collagen fiber sliding) to fiber-dominant load translation in the enthesis. Alternatively, repeated loading in growing enthesis leads to reduced mineral deposition at subchondral and apophyseal bone and disruption of ECM composition in young enthesis that functionally translate to significant decrease of enthesis toughness and tensile strength. These results are in agreement with the role of mineral as the primary toughening constituent of the enthesis (Golman et al. 2021).

The reduced mineralization and ECM disruption in loaded young calcaneal growth plate, apophysis and enthesis suggest an unexplored role of mechanical loading in enthesis and apophysis adaptation during growth. The role of increased loading on the formation and maturation of the mineralizing growth plate during postnatal maturation was elucidated in our bulk RNAseq studies, and we identified key pathways that are transcriptionally activated in the growing calcaneus as well as tendon. Activation of Rho GDP-dissociation inhibitor (RHOGDI) signaling and downregulation of mechanosensitive pathways (e.g., RAC, actin cytoskeleton) in tendon was surprising, especially considering the robust structural and morphological changes we observed in the loaded enthesis. Our findings model the structural and mechanical changes similar to those occurring in pediatric apophyseal pathologies and in the young athletes (Binks et al. 2015; Holt et al. 2020; Pennock et al. 2016; 2018). This study provides the first small animal model of pediatric apophyseal pathologies that can be leveraged for future systematic examination of mechanisms of injury and repair in growing skeleton.

This study was not without its limitations. Although muscle contraction was confirmed throughout the experimental loading duration, the translated loads from muscle to tendon, enthesis, and bone during loading are unknown. There may be unknown physiological changes that directly or indirectly affect the mechanoadaptation of the tendon and enthesis.

## Materials and Methods

All procedures were approved by the Institutional Animal Care and Use Committees at the University of Delaware and the University of Michigan. Doxycycline-inducible ACTA1-rtTA;tetO-cre mice (Acta1-Cre; C57BL6J background) and Ai32 reporter mice (C57BL6J background) were obtained from Jackson Laboratory (Bar Harbor, ME, USA). Acta1-Cre and Ai32 reporter mice were bred to generate F2 (Ai32 homozygous) lines that expressed a YFP-fused ChR2 light-sensitive opsin (455nm sensitivity) in skeletal muscle when treated with tetracycline. Dams were treated with doxycycline chow at the time of mating and maintained on chow until offspring were weaned. Offspring were genotyped using PCR (Transnetyx, Cordova, TN, USA). A total of 59 mice were used in this study. ActaCre; Ai32 homozygous mice (n=40 total) were used for unilateral light-induced stimulation of muscle in Young (4 weeks old, n = 32; 14 females and 18 males) or Adult (6-8 months old, n = 8; 2 females and 6 males) mice. Contralateral (within-animal) limbs were used as paired controls. Additionally, a second group of ActaCre; Ai32 mice was used to compare age-matched littermates as naïve controls for both Young (n = 15; 5 females and 10 males) and Adult (n = 4; 2 females and 2 males) groups.

### Optogenetics stimulation for daily tendon enthesis loading

Mice were anesthetized with isoflurane, and hair was removed using chemical hair remover from their right triceps surae muscles (Nair, Church & Dwight, Ewing, NJ, USA). Animals were placed on a heating pad to maintain body temperature at 37°C (Stoeling, Wooddale, Il, USA). The right limb was stabilized at the knee joint, and the right foot was placed on a foot pedal connected to a servomotor shaft (Aurora Scientific, Ontario, Canada) to measure isometric muscle contractions and joint torque. Animals were weighed daily before stimulations. For light stimulation, a collimated LED light (455 nm, 900 mW, M455L3, Thorlabs) and a high-power LED driver (DC2200, Thorlabs, Newton, NJ) were used for pulse modulation (Ganji et al. 2021). Driver code was adjusted in LabView (National Instruments, Austin, TX, USA) for additional Control of the pulsed light activation frequency. Triceps surae muscles were stimulated using an approximated 10Hz pulsed light (70ms on, 30ms off; 10 cycles) followed by 4 seconds of rest for 20 minutes (240 consecutive loading cycles) in Young (n = 27, 12 females and 15 males) and Adult (n = 8, 2 females and 6 males) groups. Daily bouts of loading were repeated weekly with five days of stimulation and two days of rest. A subset of these mice in the Young group (n=7) were also used to measure ankle torque throughout the stimulation using an Aurora 3-in-1 system. All other mice were loaded in a 3D printed mimic of the platform. Knee and ankle placements were replicated for the Young Control group for 20 minutes daily under anesthesia. After recovery from anesthesia, mice were returned to their cages for unrestricted cage activity. An additional cohort of Young mice were added to compare the effects of bout duration (5 minutes “short” loading; 60 consecutive cycles; n = 5, 2 females and 3 males) loading duration for 5 days.

### Micro-computed tomography

Mice were euthanized using carbon dioxide asphyxiation 48hr after the final bout of loading. The skin was carefully dissected, and distal hindlimbs (knee, ankle, and foot) were fixed for 24hr in 4% paraformaldehyde. Calcaneus-tibial complexes were scanned using micro-computed tomography (microCT) at the University of Delaware (microCT, Bruker SkyScan 1276; 59kV, 175μA, 10.58μm voxel size, 930ms exposure, 0.5mm aluminum filter). Digital reconstructions of the calcaneal apophyses were segmented based on the growth plate morphology. The posterior calcaneal tuberosity (PCT) was segmented from the 1.5mm wide posterior region of the calcaneus both in Young and Adult calcanei. Segmented regions of interest were analyzed using CTAn software (Bruker, Kontich, Belgium). Bone morphometry was assessed at the volume of interest using tissue volume, TV; bone volume, BV; bone volume/tissue, BV/TV; and bone mineral density, BMD (Campbell and Sophocleous 2014). Aligned image stacks were overlayed for qualitative assessment of the Young and Adult loaded and contralateral limbs using Dragonfly software (Object Research Systems, Montreal, Quebec).

### Histology

Distal hindlimbs from mice treated with long and short loading (20-minute and 5-minute groups) as well as from naïve controls were decalcified in 14% ethylenediaminetetraacetic acid and processed for paraffin embedding. Tissues were sectioned at 6μm in the sagittal plane and stained with Hematoxylin and Eosin (H&E) or Toluidine Blue and cover-slipped with an acrylic mounting media (Acrymount, Statlab, McKinnery, TX, USA) or used for immunohistochemistry. For immunohistochemistry, paraffin sections were deparaffinized and quenched in 0.3% hydrogen peroxide (Santa Cruz, Dallas, TX), followed by heat-mediated antigen retrieval and blocking (5% goat serum in phosphate buffer saline). Slides were then incubated in primary antibodies (listed in Table 1) overnight at 4°C, followed by washing and incubation with appropriate secondary antibodies. Horseradish peroxidase 3,3’-Diaminobenzidine system (Millipore Sigma, Billerica, MA, USA) was used for detection, slides were counterstained with Hematoxylin, and coverslipped with Acrymount (Acrymount, Statlab, McKinney, TX, USA). All slides were imaged on a brightfield microscope (Eclipse Ni-U, Nikon).

**Table 1:** Antigen retrieval and antibody information for immunohistochemistry experiments.

| Antibody | Source and Catalog # | Dilution | Buffer; Antigen Retrieval |
| --- | --- | --- | --- |
| Anti-Collagen III<br>(rabbit polyclonal) | Abcam, ab7778 | 1:100 | Tris EDTA, pH 9.0; 65°C, 2 hr |
| Anti-Aggrecan<br>(rabbit polyclonal) | Abcam, ab216965 | 1:100 | 0.1% trypsin; 37°C, 30 min |
| Biotinylated goat<br>anti-mouse IgG | Millipore-Sigma,<br>20777 | 1:1000 | Room temperature, 10 minutes |
| Biotinylated goat<br>anti-rabbit IgG | Millipore-Sigma,<br>20715 | 1:1000 | Room temperature, 10 minutes |

### Biomechanical testing of the Achilles enthesis

After 3 weeks of daily (20min) loading, Young and Adult mice were euthanized, and carcasses were stored intact at -20°C until testing. Carcasses were thawed at 4°C for no more than 24 hr and Achilles tendon-calcaneus complex was carefully dissected and stored in PBS for no more than 2 hr at room temperature. Before mechanical testing, the muscle was carefully dissected from the Achilles tendon, the plantaris tendon was detached, and the calcaneus was disarticulated from the foot. Cross-sectional area (CSA) measurements were made using photogrammetry with a custom 3D-printed fixture attached to a motor controller (Arduino, Ivrea, Italy). Forty consecutive images of the tendon-bone complex were acquired at 9-degree rotation steps using a macro lens (Basler Fujinon Lens, Ahrensburg, Germany). Images were converted from sparse to dense point clouds, and an STL surface mesh of the tendon was generated using Metashape software (Agisoft, St. Petersburg, Russia). CSA was measured using Dragonfly software at the Achilles tendon near the enthesis. The calcaneus was then placed in a custom 3D printed fixture to secure the bone (FormLabs 3B, Somerville, MA, USA), and the proximal Achilles tendon was clamped in a thin film grip (Imada, Northbrook, IL, USA).(Kurtaliaj et al. 2019) The assembled tendon-grip unit was placed in a custom PBS bath maintained at 37ºC via a temperature controller (MA160, Biomomentum Inc, Laval, Quebec, Canada) and secured with a pin to a tensile testing frame with multi-axis load cells (Mach-1 VS500CST, ±70N, Biomomentum, Laval, Quebec, Canada). Samples were preloaded to 0.01N, and gauge length (L_0_) was measured (4.01 ± 0.17 mm) as the distance between the calcaneal fixture and the thin film grip. Tendons were preconditioned for 10 cycles (±0.05N at 0.03 Hz) followed by load to failure at 0.05mm/sec. Load and torque data (6 degrees of freedom) were collected to assess off-axis loading for the duration of the experiment. Using Mach-1 Analysis and custom Matlab script (R2017 or later, Mathworks, Natick, USA), mechanical properties of the entheses were calculated from force-displacement data. Engineering stress was calculated as the instantaneous force divided by the original CSA (calculated from photogrammetry). Instantaneous grip-to-grip strain was calculated as the displacement divided by original gauge length, L_0_. Stiffness and elastic modulus were calculated from the linear portion of the load-displacement and stress-strain curves, respectively. Maximum load, maximum stress, and maximum strain were calculated, as well as work to maximum force (area under the curve, AUC) and toughness from the area under the force-displacement and stress-strain curves. Maximum strain was calculated as the strain at maximum force.

### Collagen hybridization and damage visualization

Young mice were loaded for 20 minutes daily for 5 days (n=4). Immediately following the last bout of loading, 20µl sulfo-Cy7.5 collagen hybridizing peptide (sCy7.5-CHP) (3Helix, Salt Lake City, UT, USA) was injected subcutaneously adjacent to the Achilles tendon mid-substance using an insulin syringe (Exel International, CA, USA). Mice were imaged under anesthesia (Li-Cor Pearl imager; 85 µm pixel resolution; 700 and 800 nm wavelength) 24 hours after the injection. Intensity measurements were taken in a user-defined ROI at the Achilles tendon and enthesis, using Li-Cor image acquisition software for both 700 and 800 nm wavelengths.

### RNA-sequencing of tendon and bone

Mice were euthanized 3hr after the 5^th^ or 12^th^ consecutive day of loading and tendon and calcaneus from loaded and contralateral (not loaded) limb were immediately dissected under RNase-free conditions and snap froze (n=3 male mice per time point). Age-matched, uninjured Achilles tendons were also be collected (control group; n=3 male mice). Tissues were mechanically pulverized in Trizol and total RNA was isolated using spin-columns with DNA digestion. poly-A mRNA libraries were prepared and next-generation sequencing was performed using Illumina NovaSeq Shared platform with Snakemake pipeline analysis for quality control and sequence alignment (X. Zhang and Jonassen 2020). Differential gene expression was determined from count matrices with either paired design (loaded vs. contralateral) or unpaired design (loaded or contralateral vs. control) in DESeq2 in R/Bioconductor (Love, Huber, and Anders 2014).

### Statistical analysis

Statistical comparisons were performed using Prism (8.3 or higher, GraphPad Software, La Jolla, USA) or in R. Gaussian distribution of the data was evaluated using Shapiro-Walik and D’Agostino-Pearson normality tests. Results from micro-CT analyses were tested for normality and compared between the loaded and control samples using an unpaired t-test. For statistical comparison of the CSA measurements and mechanical testing results, parametric data were tested using paired two-tailed t-test, and Wilcoxon rank test was used for non-parametric results. For RNAseq data, we used Wald tests of significance and p-values to correct for multiple testing using the Benjamini and Hochberg method (p-adjusted <0.05). For pathway analyses and identifying upstream mechano-sensitive regulators, paired and unpaired gene lists were analyzed using Ingenuity Pathway Analysis (Qiagen) with an input cut-off of p-adjusted < 0.1. After IPA analysis, we filtered with two significance parameters to include only enrichment |z| > 2 and Fisher’s exact p < 0.05 and <0.01 for pathways and regulators, respectively. For power analysis, the effect size was calculated from the descriptive statistics of each group, and the effect size was calculated as the ratio of mean and standard deviation (SD) of difference. Power for each variable was calculated using total sample size (N=16), significance (α=0.05), and the calculated effect size for a t-test with match mean values. For power < 0.8, a priori power analysis was performed to calculate the required sample size to achieve 0.8 power. For most outcomes, statistical significance was considered as *p* = or < 0.05. Mean ± SD are reported.

